# Aneuploidy influences the gene expression profiles in *Saccharomyces pastorianus* group I and II strains during fermentation

**DOI:** 10.1101/2021.12.09.471776

**Authors:** Roberto de la Cerda, Karsten Hokamp, Fiona Roche, Georgia Thompson, Soukaina Timouma, Daniela Delneri, Ursula Bond

## Abstract

The lager yeasts, *Saccharomyces pastorianus*, are hybrids of *Saccharomyces cerevisiae* and *Saccharomyces eubayanus* and are divided into two broad groups, Group I and II. The two groups evolved from at least one common hybridisation event but have subsequently diverged with Group I strains losing many *S. cerevisiae* chromosomes while the Group II strains retain both sub-genomes. The complex genomes, containing orthologous alleles from the parental chromosomes, pose interesting questions regarding gene regulation and its impact on the fermentation properties of the strains. Superimposed on the presence of orthologous alleles are complexities of gene dosage due to the aneuploid nature of the genomes. We examined the contribution of the *S. cerevisiae* and *S. eubayanus* alleles to the gene expression patterns of Group I and II strains during fermentation. We show that the relative expression of *S. cerevisiae* and *S. eubayanus* orthologues is positively correlated with gene copy number. Despite the reduced *S. cerevisiae* content in the Group I strain, *S. cerevisiae* orthologues contribute to biochemical pathways upregulated during fermentation which may explain the retention of specific chromosomes in the strain. Conversely, *S. eubayanus* genes are significantly overrepresented in the upregulated gene pool in the Group II strain. Comparison of the transcription profiles of Group I and II strains during fermentation identified both common and unique gene expression patterns, with gene copy number being a dominant contributory factor. Thus, the aneuploid genomes create complex patterns of gene expression during fermentation with gene dosage playing a crucial role both within and between strains.

## Introduction

The yeasts used in lager beer production have long been recognised as being unique and distinct from those used for making ales (1). In 1870, in recognition of their unique physiological qualities, this group of yeast were given a taxonomical classification of *Saccharomyces pastorianus* by Max Reess (2). After more than a century of genetic studies, we now know that *S. pastorianus* strains are natural hybrids of *Saccharomyces cerevisiae* and *Saccharomyces eubayanus* (3, 4). While *S. cerevisiae* has long been associated with beer production, especially for ales, *S. eubayanus* was only discovered in Patagonia, South America, in 2011 (5). While Patagonia is considered to be the primary radiation source of *S. eubayanus*, natural isolates have also been found in North America, China, Tibet and New Zealand but to date, no isolates have been identified in Europe (6–9) with only limited metagenomic evidence for the presence of this species in Italian Alps (10). The hybridisation events that generated the current strains of *S. pastorianus* are estimated to have occurred some 500-600 years ago following the introduction of *S. eubayanus* into Central Europe from China or Tibet, most likely along trade routes such as the Silk Road. An inherited property of cryotolerance from the *S. eubayanus* genome allows *S. pastorianus* strains to ferment at temperatures as low as 7-13°C (11). This property together with the robust fermentation kinetics inherited from the *S. cerevisiae* genome created yeast hybrid strains capable of producing a crisp clean tasting lager that is now the most favoured alcohol-containing beverage.

Genome analysis of *S. pastorianus* strains identified two distinct types, based on genome content and chromosome composition (4, 12–17). Group I, or Saaz strains, are typically triploid in DNA content, retaining all the parental *S. eubayanus* chromosomes but have lost many *S. cerevisiae* chromosomes. This group includes several strains isolated from breweries in Bavaria (Germany) and Bohemia (Czechia) and includes the strains CBS1538, CBS1513, and CBS1503 that were originally isolated in The Carlsberg Laboratory in the late 19^th^ century (18). The Group II, or Frohberg strains, include isolates from Dutch, Danish and North American breweries and are mainly tetraploid in DNA content, containing approximately 2n *S. cerevisiae* and 2n *S. eubayanus* genome content. Both groups display chromosomal aneuploidy with chromosome numbers ranging from one to six (19, 20). In addition to the parental chromosomes, *S. pastorianus* strains contain several hybrid chromosomes containing both *S. cerevisiae* and *S. eubayanus* genes that resulted from recombination, at precise locations, between the parental chromosomes. Some of the recombination breakpoints are located within coding regions, creating a set of hybrid genes unique to lager yeasts (4, 12, 21, 22).

Based on genome analysis, the current understanding of the origin and evolution of *S. pastorianus* strains is that both Group I and II strains evolved from a common hybridisation event between *S. eubayanus* and *S. cerevisiae* strains to generate a progenitor hybrid. Subsequently, this progenitor strain underwent a second hybridisation event with a second *S. cerevisiae* strain to generate Group II strains (4, 23). This model is supported by shared recombination sites on hybrid chromosomes, evidence of Single Nucleotide Polymorphisms (SNPs) in the *S. cerevisiae* genome of Group II strains and differences in telomeric regions in Group I and II strains (23). The genome data is also consistent with a scenario in which both groups emerged from a single hybridisation event between a diploid *S. eubayanus* strain and a heterozygous *S. cerevisiae* diploid strain, with Group I strains experiencing a selective loss of a significant proportion of the heterogeneous *S. cerevisiae* genome (16, 20, 24). Both groups then evolved independently with each undergoing further recombination events between the sub-genomes (3, 20, 22, 24). Both groups have subsequently diverged to create distinct sub-groups, each with their own unique physiological and biological properties. Fermentation analysis of Group I and II strains reveals that each group produces distinctive aroma and flavour profiles (25).

The complex genome of *S. pastorianus*, containing orthologous alleles emanating from different parental chromosomes, poses interesting questions regarding gene regulation and its impact on the fermentation properties of the strains. Superimposed on the presence of orthologous alleles are complexities of gene dosage due to the aneuploid nature of the genomes. Aneuploidy has been shown to influence gene expression patterns in eukaryotic cells (26–29). The presence of gene orthologues from two different parental *Saccharomyces* species, together with copy number differences between the orthologues has the potential to affect the cellular proteome and specifically the stoichiometry of *S. cerevisiae* and *S. eubayanus* proteins within protein complexes (30). Furthermore, gene copy number differences between the Group I and Group II strains may lead to differences in cellular physiology and thus the fermentation properties of the two types of *S. pastorianus*.

Previous transcriptome analyses of *S. pastorianus* strains under fermentation conditions were limited by technology and mainly focussed on the analysis of *S. cerevisiae* genes (31–34). Such studies predated the discovery of *S. eubayanus* as a contributing parent to *S. pastorianus* and advances in RNA sequencing technologies (35). More recent transcriptome studies focussed on the analysis of sub-sets of genes with specific roles in fermentation or in specific physiological conditions such as cold storage or responses to temperature (36–38). Important findings regarding the gene expression of genes involved in maltose utilisation, carbohydrate metabolism, glycerol mobilisation, anaerobiosis, and protein biosynthesis have emerged from these studies, providing a road map for a more detailed transcriptome analysis of these industrial strains.

Here, we analysed the transcriptomes of the Group I, CBS1538 and the Group II, WS 34/70 *S. pastorianus* strains and specifically examined the contribution of the two sub-genomes to the gene expression patterns under fermentations conditions. Using *de novo* genome sequencing of these strains, we related gene expression patterns to gene copy numbers. We show that the relative expression of *S. cerevisiae* and *S. eubayanus* orthologues is directly correlated to gene copy number in the Group II strain. Gene copy number plays a smaller role in gene expression patterns in the Group I strain, most likely due to the more limited genetic redundancy. Despite the reduced *S. cerevisiae* content in the Group I strain, *S. cerevisiae* orthologues contribute significantly to biochemical pathways upregulated during fermentation, specifically to amino acid metabolism. Finally, comparison of the transcript patterns in the Group I and II strains identified both common and unique gene expression patterns during fermentation.

## Results

### Fermentation profiles of Group I and II strains

Two *S. pastorianus* strains, CBS1538 and WS 34/70, representative of Group I and Group II strains respectively, were chosen for analysis. Both strains displayed similar fermentation profiles in 10% wort up to Day 4 but thereafter, strain WS 34/70 fermented faster than strain CBS1538 and reached a lower final attenuation (Figure 1A). At the end of fermentations in 3L tall tubes, the two strains produced similar volatile compound profiles with the exception that ethyl butyrate and methionol levels were higher in WS 34/70, and ethyl hexanoate was produced in higher levels in strain CBS1538 (Figure 1B). Similar volatile profiles were obtained in small scale (15 mL) fermentations, the scale used for RNA extractions, with the exception that higher levels of acetaldehyde were observed in both strains (data not shown).

**Figure 1.**
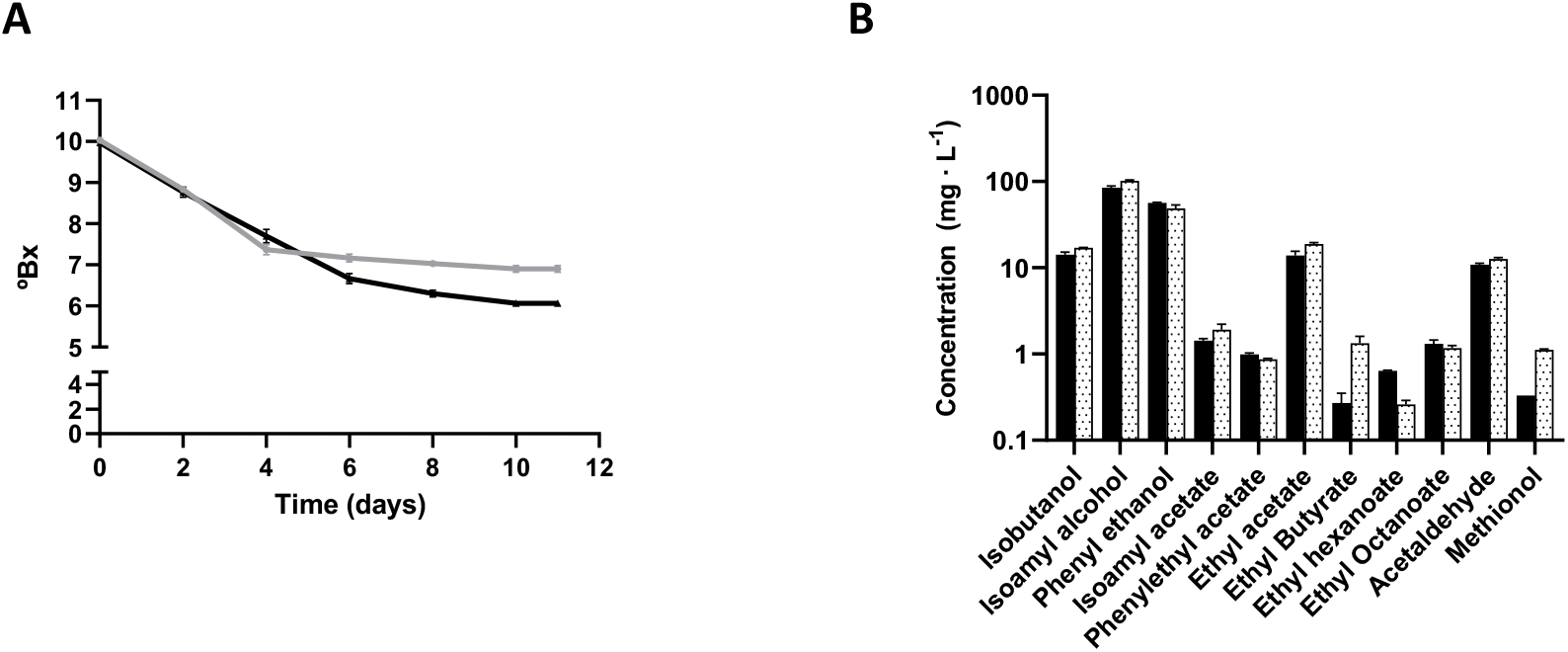
Fermentation profiles of the Group I CBS1538 and Group II WS34/70 strains. **A.** Fermentations were carried out in 10%Wort at 13°C in 15 mL cylindrical tubes. Sugar consumption was measured with a Brixometer. CBS1538: grey line, WS34/70, black line. Error bars represent the standard deviations from the mean of triplicate fermentations. **B.** Volatile profiles of compounds present in wort at the end of fermentations. Fermentations were carried out in 10% Wort at 13°C in 3L tall tubes. Error bars represent the standard deviations from the mean of duplicate fermentations.

### Chromosome composition of strains CBS1538 and WS 34/70

To compare the transcriptomes of Group I and Group II lager yeasts, RNA was extracted from strains CBS1538 and WS 34/70 on Day 2 and Day 4 of small-scale fermentations in 10% wort. These time points were chosen as previous studies have shown that this is the period of maximum metabolism during the fermentation (32). RNA was also extracted from the same strains grown in minimal medium without amino acids to provide a baseline for comparison. To map the transcripts, the genomes of CBS1538 and WS 34/70 isolates were sequenced *de novo* and annotated using data from the annotated and fully assembled reference genome *S. pastorianus* 1483 (Group II strain) as well as to a combined genome assembled from the parental reference genomes *S. cerevisiae* and *S. eubayanus*. Both approaches yielded highly similar results, however since the *S. pastorianus* 1483 genome lacked some information for *S. cerevisiae* genes on chromosomes III and VII, the data from mapping to the combined parental genomes was used for this analysis. Information from genome sequencing and mapping of the two strains (Figure 2A, B) confirmed the absence of *S. cerevisiae* chromosomes II, III, IV, IV, VII, VIII, XII, XIII, XV and XVI in CBS1538 (Figure 2A). Furthermore, we observed alterations in the copy number of chromosomes and differences in hybrid chromosomes to the previously reported chromosome content of both strains (16, 22, 24) (Figure 2C). The estimated chromosome copy numbers were 48 and 76 for CBS1538 and WS 34/70 respectively.

**Figure 2.**
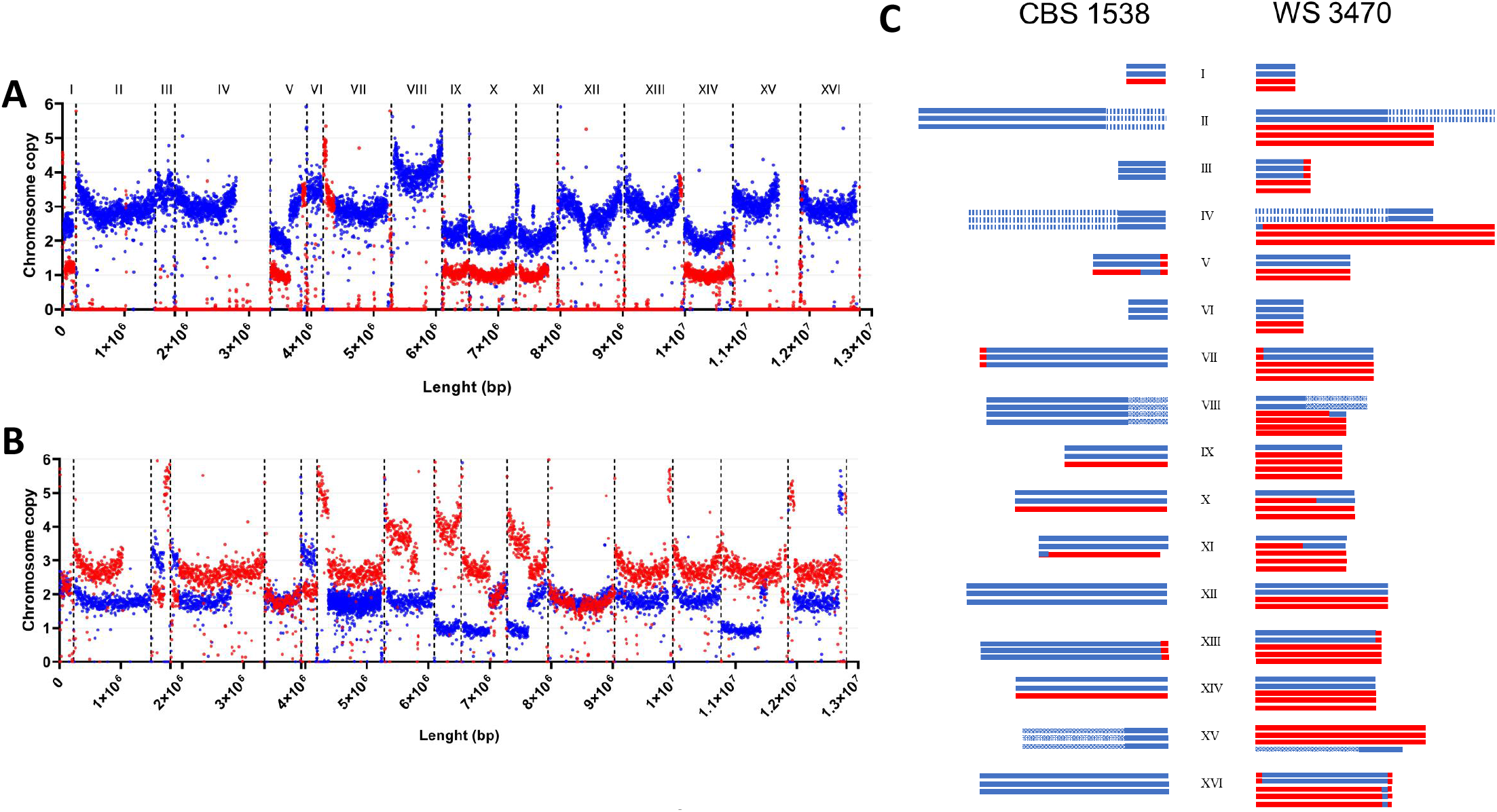
Chromosome copy number of CBS1538 (**A**) and WS 34/70 (**B**) genomes. Estimated copy number and chromosome types for *S. eubayanus* (blue) and *S. cerevisiae* (red) isolates from *de novo* sequencing mapped to the combined parental genomes. The start of each chromosome is shown by a vertical dotted line. *, denotes translocations between *S. eubayanus* chromosomes II/IV, IV/II, VIII/XV, XV/VIII present in both strains. Hybrid chromosomes are evident from the change in copy number within a chromosome. **C**. estimated types and copy number of *S. cerevisiae* (red) and *S. eubayanus* (blue) and hybrid (red/blue) chromosomes in CBS1538 and WS 34/70.

### Gene expression profiles of Group I and II strains during fermentation

Transcripts were either assigned as *S. cerevisiae* (Sc) or *S. eubayanus* (Se) based on sequence identity and chromosome assignment. A total of 11335 transcripts/Open Reading Frames (ORFs) were detected for the WS 34/70 strain of which 5083 were assigned as *S. eub*ayanus and 6252 as *S. cerevisiae* (ratio Se:Sc = 0.81:1) (Supplemental Table 1). For the Group I strain, a total of 6798 transcripts were detected, 5163 were *S. eubayanus* and 1635 were *S. cerevisiae* (ratio Se:Sc = 3.16:1). These values agree well with the previously determined gene count for the two strains (39). We examined the differential gene expression (DEG) of transcripts mapped to each strain in three different experimental conditions (minimal medium without amino acids, wort Day 2 and wort Day 4) (Supplemental Tables 2-4). In the WS 34/70 strain, the greatest number of genes displaying DEG was detected in the comparison of cells grown in minimal medium and wort on Day 2 where 42% of transcripts were differentially expressed, while 28% of genes were differentially expressed between growth in minimal medium and in wort on Day 4. A much lower level of DEG was detected between cells grown in wort on Days 2 and 4 where just 14% of genes were differentially expressed. The pattern of DEG in CBS1538 was similar to that observed with the WS 34/70 strain with 32% of transcripts differentially expressed between growth in minimal medium and wort on Day 2, 23% between growth in minimal medium and wort on Day 4 and 16% between growth in wort on Days 2 and 4.

As *S. pastorianus* contains both *S. cerevisiae* and *S. eubayanus* sub-genomes, we examined the DEG patterns for *S. cerevisiae* or *S. eubayanus* alleles under the three different experimental conditions. In the WS 34/70 strain, *S. eubayanus* genes are significantly over-represented in the differentially upregulated gene pool under all three conditions but were expressed at the expected ratio in the differentially downregulated gene pool (Figure 3A). Interestingly, on Day 2 relative to Day 4 in wort, there are twice as many Se alleles upregulated compared to Sc alleles. In the CBS1538 strain, Se alleles are overrepresented in both the upregulated and downregulated gene pools in cells in wort on Day 2 relative to Day 4 and in wort on Day 2 relative to minimal medium and in the downregulated gene pool in wort on Day 4 relative to minimal medium.

**Figure 3.**
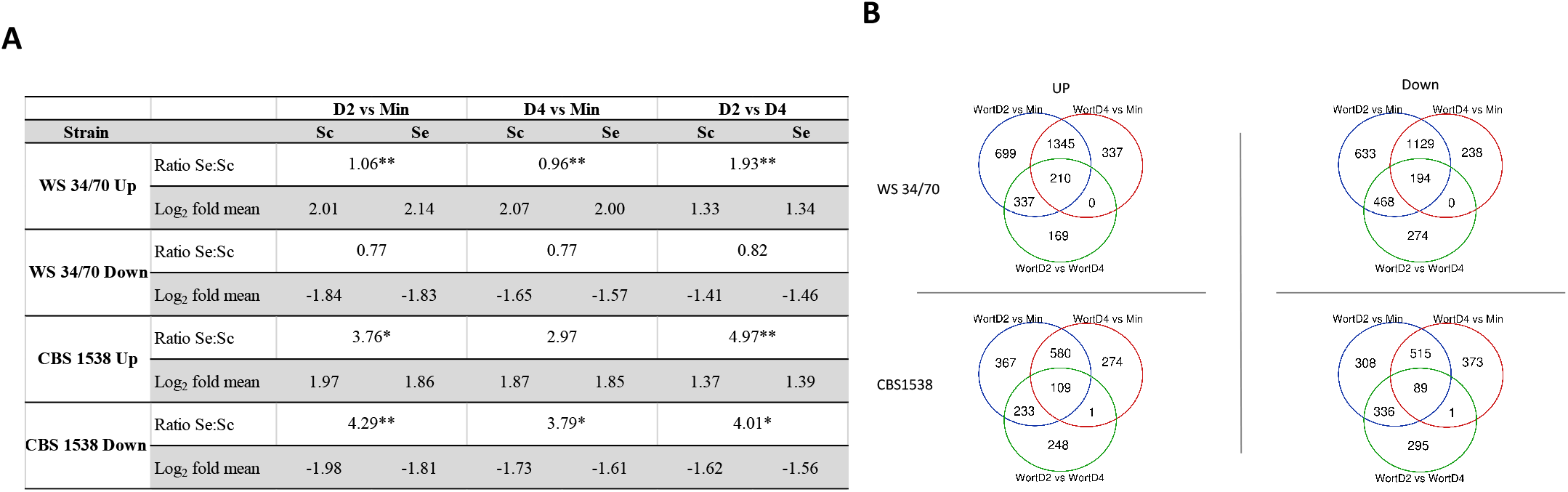
**A.** Ratio of *S. cerevisiae* (Sc) and *S. eubayanus* (Se) in differentially up-regulated and down-regulated transcript pools (log_2_-fold change ≥1 or ≤ −1) and mean log_2_-fold changes under the three conditions in CBS1538 and WS34/70. Ratios of differentially expressed Se:Sc genes with significant differences from the expected are shown with an asterisk, p≤0.001**, p≤0.05*. **B.** Venn diagrams showing the relationship of differentially up-regulated and down-regulated transcript pools under the three conditions shown.

The mean log_2_-fold change for *S. cerevisiae* and *S. eubayanus* alleles in both the WS 34/70 and CBS1538 strains are extremely consistent under all conditions indicating that both Group I and Group II strains, and their Se and Sc sub-genomes, respond similarly to the physiological conditions. The mean log_2_-fold change and the range of DEG on Days 2 and 4 wort was less than that observed for differential expression between minimum medium and wort on Days 2 and 4 for both strains with the exception for down regulated genes on Days 2 and 4 in CBS1538 (Figure 3A and Supplemental Fig1).

We next examined the relationship of DEG under the three conditions in the two strains. As might be expected, a large number of genes are differentially expressed between growth in minimal medium and in wort on Days 2 and 4 in both the Group I and Group II strains (Figure 3B). While fewer in number, there is an overlap in the genes differentially expressed in wort Day 2 relative to Day 4 and in wort Day 2 relative to minimal medium. On the other hand, there are no genes differentially expressed that are specific to wort Day 2 relative to Day 4 and to wort Day 4 relative to minimal medium in WS 34/70 and just 1 gene in this category in CBS1538.

Only a small number of genes are commonly upregulated or downregulated respectively in all three conditions in the Group I and the Group II strains (Figure 3B, Supplemental Table 5). The commonly upregulated gene pool in WS 34/70 (n=210), contains 23 pairs of Sc and Se orthologous alleles, including, *ADH1*, *ADH5*, *PGK1*, *GPH1*, *HXK1*, *ENO1*, that are central to carbohydrate metabolism and *ILV6*, required for branched chain amino acid biosynthesis. Likewise, both Sc and Se alleles of *LEU4*, *LYS1* and *SPD1* are upregulated in all three conditions in CBS1538. Orthologous gene pairs are also observed in the commonly downregulated gene in both strains (Supplemental Table 5).

### Gene ontologies enriched in Group I and II strains under fermentation conditions

Gene ontology analysis of the differentially expressed genes under the three conditions in the Group I and Group II strains revealed an enrichment in both carbohydrate and amino acid metabolism on Day 2 of fermentation in the upregulated gene pool (Figure 4A). Genes required for the utilisation of all major sugars including sucrose, fructose, galactose as well as pentose sugars along with the genes required for glycolysis and pyruvate metabolism are all upregulated in wort Day 2 relative to minimum medium and a subset of these genes are upregulated on Day 2 relative to Day 4 and on Day 4 relative to minimum medium in the Group II strain. Except for genes involved in starch and sucrose metabolism, the observed gene set for carbohydrate metabolism in the Group II strain were not enriched in the Group I strain (Figure 4A). Genes for the biosynthesis and metabolism of amino acids are enriched during the fermentation in wort in both Group I and Group II strains (Figure 4A). In addition to the enrichment of genes for these major metabolic activities in the two strains, we observed differences in specific gene ontologies between the two strains, with genes involved in glycerolipid metabolism, porphyrin metabolism, ABC transporters, peroxisome activity and longevity upregulated in the Group II strain and not the Group I strain.

**Figure 4.**
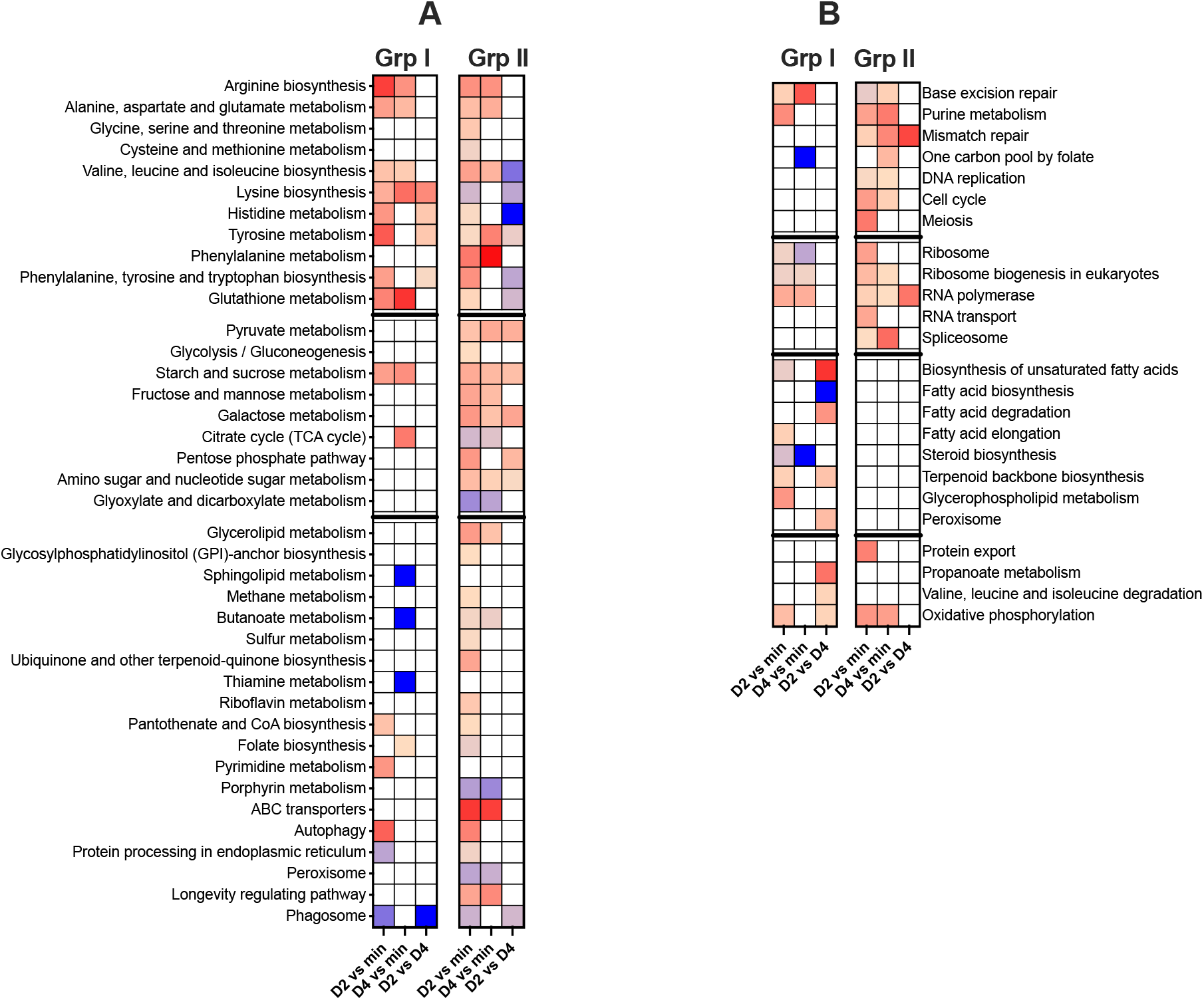
Gene ontology analysis of differentially expressed genes in Group I and Group II strains. Genes displaying a log_2_-fold change ≥1 **(A)** or ≤ −1 **(B)** were examined for enriched gene ontologies using Cluego. Graph shows deviations from the expected ratio of Sc to Se alleles; expected ratio is buff, >expected ratio; buff to red gradient, < expected ratio; buff to blue gradient. Blank cells, no enrichment. Group I expected ratio Sc:Se =0.24:0.76, Group II, Sc:Se = 0.55:0.45.

Genes involved in nucleic acid metabolism, DNA repair and DNA replication are downregulated under fermentation conditions and on Day 2 relative to Day 4 in wort in the Group II strain with a subset of these genes similarly regulated in the Group I strain (Figure 4B). Likewise, genes involved in major anabolic pathways such as protein synthesis including genes required for ribosome biogenesis and rRNA, mRNA and tRNA processing are downregulated in wort and on Day 2 relative to Day 4 in both strains. Interestingly, a set of genes involved in fatty acid, glycerophospholipid and steroid metabolism, required for the production of membrane components, are uniquely downregulated in the Group I strain (Figure 4B).

We examined the contribution of the Sc and Se sub-genomes to these enriched pathways. Se and Sc alleles were observed to contribute at the expected Se:Sc ratio or in a higher proportion than expected to the differentially regulated gene pool (Figure 4A, B). Se alleles contribute in a higher proportion than expected on Days 2 and 4 of fermentation in the Group II strain (Figure 4A). Conversely, we observed that Sc alleles contribute in a greater proportion than expected to amino acid metabolism in the Group I strain under all three conditions (Figure 4A). For some specific pathways, we observed that Se alleles accounted for all the differentially regulated genes, for example Se alleles are used exclusively in histidine metabolism on Day 2 relative to Day 4 in the Group II strain and to phagosome associated genes in wort on Day 2 relative to Day 4 and to sphingolipid, butanoate and thiamine metabolism on Day 4 relative to minimum medium and to phagosome activity on Day 2 relative to Day 4 in wort in the Group I strain (Figure 4A).

For the downregulated genes, we observe that Sc alleles contribute to gene ontologies at the expected ratio, and in some pathways, in a higher proportion than expected to the ratio of Sc:Se for the whole genome in both strains while Se alleles exclusively contribute to three categories, namely one carbon pool by folate, fatty acid and steroid biosynthesis in the Group I strain (Figure 4B).

### The effect of gene dosage on gene expression profiles

As *S. pastorianus* strains are aneuploid in nature and have different copy numbers of *S. cerevisiae* and *S. eubayanus* chromosomes, we were interested to determine if gene expression profiles were influenced by the copy number of Sc and Se alleles. The copy number for each Sc and Se gene was calculated from the sequence coverage from the *de novo* genome sequencing. The DEG of Sc and Se orthologues within each strain was determined for the three experimental conditions and was correlated to the Sc:Se gene copy number ratio. The data for Day 2 vs min is shown in Figure 5. Sc and Se orthologues are present in varying ratios in the Group II strain (Figure 5A). The analysis of some 1555 Sc and Se orthologues indicates that the DEG of Sc and Se alleles was positively correlated to the ratio of the copy number of Sc:Se alleles (Figure 5A, B, p<0.0001, R^2^ = 0.90). As the Sc copy number increases, so too does the levels of Sc transcripts and vice versa for Se alleles. At an Sc:Se ratio of 1:2 and 3:4, Se transcripts predominate while at ratios, 3:1, 4:1 and 5:1 Sc alleles predominate. Interestingly, at a 1:1 ratio, Se alleles predominate in the Group II strain while at 3:2, 2:1 there is almost equal levels of both transcripts. The relationship between the DEG of Sc and Se orthologues and the ratio of gene copy number is observed for genes on both parental and hybrid chromosomes as exemplified in Figure 5C for chromosome X. The WS 34/70 strain has four copies of chromosome X, one Sc, one Se and 2 hybrid chromosomes (Sc/Se) with the recombination point occurring at the gene *THD2* (YJR009C) (Figure 2C). The ratio of Sc:Se to the left of *TDH2* is 3:1 and to the right 1:1 (Figure 2C). For genes to the left of *TDH2*, expression of Sc alleles predominates due to the higher Sc copy number while to the right, Se alleles predominate despite the equal copy number (Figure 5C). The correlation of DEG patterns to orthologue gene copy number ratio was observed in all three experimental conditions (data not shown).

**Figure 5.**
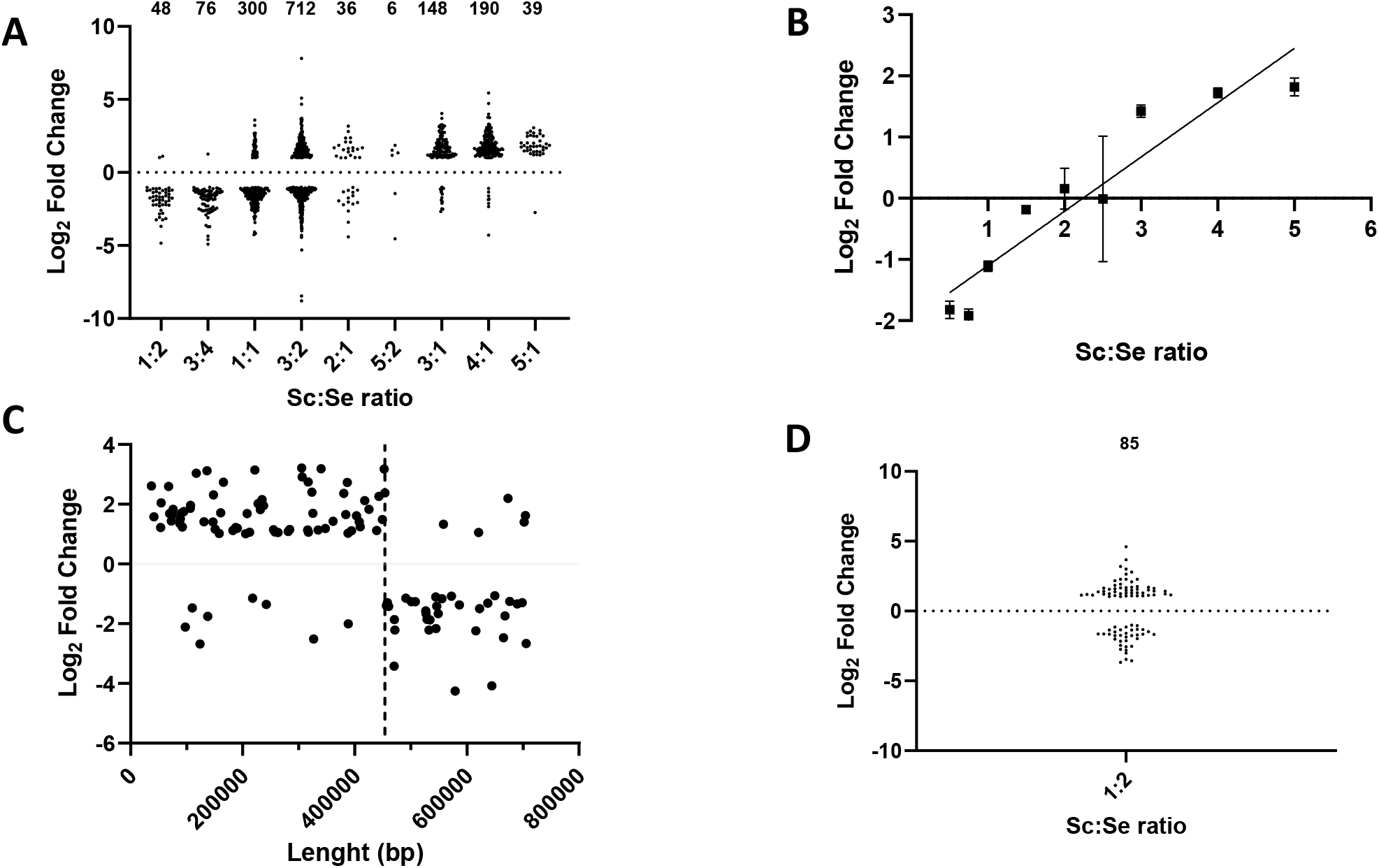
Effect of gene copy number on gene expression patterns. The DEG of Sc and Se orthologues were grouped according to the ratio of gene copy number in WS34/70 (**A**) and (**D**) CBS1538. Data for Day 2 in wort relative to minimum medium is shown. The number of paired orthologues at each copy number ratio is shown above the graphs. **B**. Correlation of differential expression of Sc:Se gene orthologues in WS34/70 with gene orthologue copy number ratio. The mean of log_2_ fold changes in expression between Sc and Se orthologues is plotted against gene orthologue copy number ratio. Error bars represent the standard error of the mean. **C.** Log_2_-fold gene expression difference in Sc and Se orthologues on chromosome X in WS34/70. The dotted line marks the recombination site (*THD2*) on hybrid chromosomes. Sc:Se ratio to left of vertical dotted line is 3:1 and after 1:1.

The analysis of the gene expression of Sc and Se orthologues in the Group I strain presented a different scenario. Firstly, due to the reduction of the Sc genome content in the Group I strain, the number of orthologues is much smaller (n=237). Secondly, the variations in copy numbers are much smaller, with most orthologues present in a ratio of Sc:Se, 1:2 (Figure 5D). There are just two alleles present in a 2:3 and one at a 2:1 ratio (data not shown). Surprisingly, we see a different expression pattern to what was observed in Group II; at the Sc:Se 1:2 ratio, a majority of Sc alleles display higher transcript levels than the orthologous Se alleles, although the reverse expression pattern is evident for some orthologues.

### Comparison of the transcriptomes of Group I and Group II strains

To directly compare the steady state mRNA landscapes in the Group I and II strains, we generated a consolidated transcriptome for each strain by obtaining the sum of Sc and Se allele transcripts encoded for each gene (Supplemental Table 6). We compared the expression of this gene set in the Group I and II strains under the three different experimental conditions (Figure 6 and Supplemental Table 7). There is a high degree of overlap between the differentially expressed gene pool in both strains suggesting that both strains respond similarly to the environmental conditions (Figure 6A). Growth in minimal medium invoked the greatest condition specific DEG between the two strains but a substantial number of genes were also specifically expressed in wort on Day 2 and Day 4 respectively. We identified 310 Group I-specific genes that are upregulated in all three conditions and likewise 299 Group II-specific genes. The list of condition specific and Group-specific genes is shown in Supplemental Table 8. An analysis of gene ontologies associated with the differentially expressed gene set reveals differences between the Group I and II strains (Figure 6B). Firstly, we observed that genes associated with ribosome biosynthesis are enriched on Day 2 and 4 in wort in the Group II strains while genes associated with metabolism of certain amino acid, pentose phosphate pathway and oxidative phosphorylation are also upregulated on Day 2 in this strain. Genes associated with DNA repair and protein processing and transport are upregulated in minimal medium. Conversely, genes associated with membrane biosynthesis, sugar metabolism and amino acid metabolism are upregulated in minimal medium in the Group I strain, and a subset of these genes are also upregulated on Day 4 in wort. Surprisingly, there are no gene ontologies enriched on Day 2 in the Group I strain (Figure 6B).

**Figure 6.**
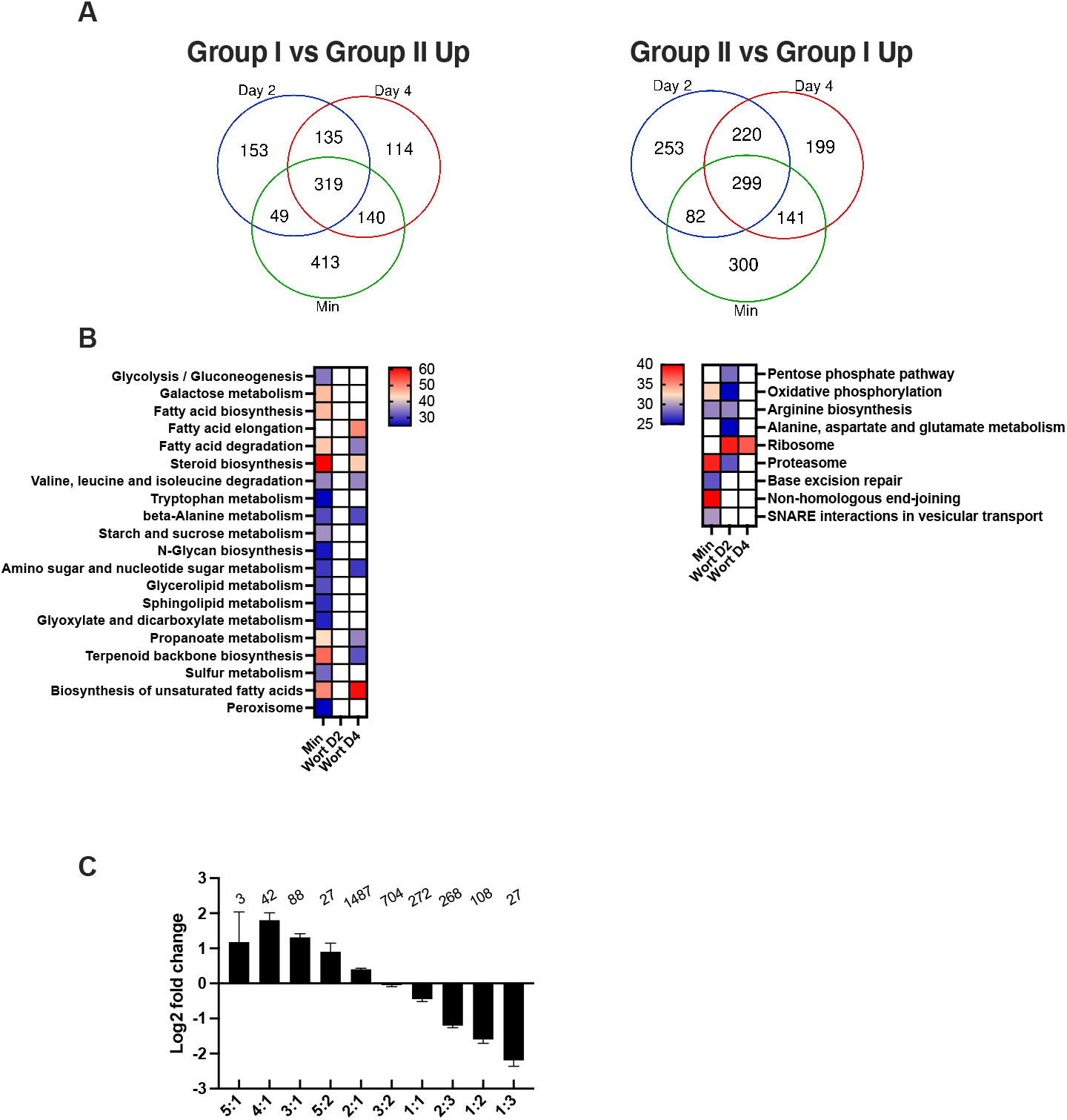
Comparison of gene expression in Group I and Group II strains. Transcript reads for Sc and Se alleles were summed in CBS1538 and WS34/70 respectively to create a combined transcriptome. Gene expression profiles in the three conditions, growth in minimal medium, in wort on Day 2 and in wort on Day 4 were compared between the two strains. The data for Day 2 in wort are shown. **A.** Venn diagram of genes upregulated in the Group I and Group II strains. **B.** Gene ontologies enriched in the Group I and Group II strains respectively. White squares, no enrichment, enriched ontologies with the shading represents the percentage of associated genes for each ontology group with the minimum set at 25%. **C.** Log_2_-fold differences in gene expression between Group I and II strains were grouped according to the ratio of Sc:Se gene copy number. The number of genes in each ratio category is shown above the graphs. Data for Day 2 in wort is shown. The error bars represent the standard error of the mean Log_2_-fold change in gene expression between Group I and II strains.

As the gene copy number for the sum of gene orthologue transcripts still varied between Group I and II strains and was shown to influence gene expression profiles within a strain, we looked to see if gene copy number influences the DEG observed between the Group I and Group II strain. To do this, we compared the log_2_ fold changes in gene expression between the two strains to the ratio of the total gene copy number (sum of all Se and Sc alleles) for each gene. The data for gene expression on Day 2 in wort is shown in Figure 6C. There is a significant positive correlation between log_2_ fold change and ratio of total gene copy number between the two strains (p,<0.005, R^2^ = 0.8). This correlation was observed across all three experimental conditions (data not shown). Thus, as observed for gene expression within a strain, gene dosage plays a significant role in the DEG between the Group I and Group II strains.

## Discussion

The origin and evolution of lager yeasts are widely debated, and several hypotheses have been proposed to account for the divergence in chromosome composition and copy number between the Group I and II strains (20, 23, 24). The major distinction between the two groups is the loss a significant portion of the *S. cerevisiae* sub-genome through the loss of whole chromosomes in the Group I strains. In addition, differences in chromosome composition, copy number and the number and type of hybrid chromosomes exist between strains within each group. Despite these differences, strains from both groups ferment sugars and produce aromatic beers, with individual strains displaying variations in the final aromatic volatile profile (25). A major difference between Group I and II strains lies in the ability to ferment maltotriose. While all Group II strains can uptake and ferment maltotriose, only a subset of Group I strains have the necessary transporters to import this trisaccharide (40–42). The Group I strain used in this study, CBS1538, does not ferment maltotriose. We confirm here the superior fermentation rates of the Group II strains and show that the two strains produce unique flavour profiles. The two strains consume sugars at the same rate up to Day 4 and thereafter the Group II strain fermented at a faster rate. WS 34/70 produces more isoamyl acetate than CBS1538, surpassing olfactory thresholds. This ester comes from the esterification of the higher alcohol isoamyl alcohol and imparts banana-like aromas, a desirable compound in beer. The WS34/70 also produced higher levels of ethyl butyrate while in contrast, CBS1538 overproduces ethyl hexanoate. Both esters contribute to the tropical fruit aroma in beer (43, 44).

We were interested in understanding how the two sub-genomes, present in the strains, contribute to the overall transcriptome during fermentation and furthermore what are the consequences for the reduction in the *S. cerevisiae* sub-genome to the gene expression patterns between the Group I and II strains. The complex genomes of lager yeasts pose several challenges to the analysis of the transcriptomes of these strains. At present, just one *S. pastorianus* Group II genome, that of strain CBS1483, has been fully annotated and assembled into chromosomes and this serves as the reference genome for *S. pastorianus* at NCBI (20), although up to 16 annotated genomes are also available https://www.ncbi.nlm.nih.gov/genome/browse/#!/eukaryotes/342/. Due to differences in chromosome copy numbers, and chromosomal rearrangements, the reference genome may not be the ideal genome for transcript mapping, for example, it differs from the two strains, WS 34/70 and CBS 1538, used in this study, as it lacks hybrid chromosomes III and VII. There are also chromosome copy number differences between the strains used here and the reference strain.

To avoid such issues, we re-sequenced the WS 34/70 and CBS1538 strains and used the combination of the parental strains *S. cerevisiae* and *S. eubayanus* genomes to map the transcriptomes. The copy number for each annotated gene was determined from the sequence coverage depth from *de novo* sequencing.

We noted some differences in the chromosome composition of both WS 34/70 and CBS1538 to the published data (16, 24, 45). Specifically, we noted one extra copy of chromosomes Sc III, VII, XIV, and XVI in relation to the most recently published WS 34/70 sequence (24).

Additionally, we reconfirm the presence of a hybrid chromosome, ScVIII/SeVIII, recombining at YHR165C, that was previously identified in the WS 34/70 strain and in another Group II strain 7012 by our group (12, 21) but which is not documented in other published sequences (16, 24). For the CBS1538, we noted just two copy number differences, an extra copy of both Se VI and SeVIII. Chromosome copy number differences are also noted between published genomes for WS 34/70 (16, 24, 45). The differences in copy numbers between studies may arise from differences in methods used to determine sequence coverage depth or may reflect genuine differences between strain isolates. The lager yeast strains emerged just some 500-600 years ago and thus may still be experiencing genomic flux in this early stage of evolution. We previously showed that the genomes of lager yeasts are dynamic and can undergo chromosome rearrangements to produce new hybrid chromosomes as well as chromosome copy number changes following exposure to stress such as high temperatures (46). Furthermore, fermentations carried out in high specific gravity wort (22°P) at an ambient temperature of 20°C, which is higher than that used for routine fermentations (13°C), led to chromosome copy number changes in a single round of fermentation (46). Thus, differences in propagation and culturing of *S. pastorianus* strains may contribute to the differences observed between strain isolates.

The analysis of the gene expression patterns reveals that the Group I and Group II strains respond similarly to the physiological conditions imposed with both strains showing similar mean log_2_ fold changes under all conditions. Interestingly, we observed that the Sc and Se sub-genomes are differentially utilised under the different physiological conditions with Se alleles contributing significantly to gene expression on Day 2 of fermentation in both strains and on Day 4 in the Group II strain.

Gene ontology analysis revealed the upregulation of pathways associated with carbohydrate and amino acid metabolism on Day 2 of fermentation although there was less enrichment of genes associated with sugar metabolism in the Group I strain. The upregulation of carbohydrate and amino acid metabolism in the early stages of fermentation is consistent with what is known about the overall metabolic activity of yeast during fermentation. Cells undergo 1-2 doublings during the first three days of fermentation and thereafter, cell numbers remain unchanged or slightly decrease. The lack of representation of genes associated with carbohydrate metabolism in the Group I strain, except for those associated with starch and sucrose metabolism, distinguishes the two strains and appears to reflect differences in metabolic activity. As overall sugar consumption is similar in the two strains on Day 2, it is possible genes associated with carbohydrate metabolism in the Group I strains are induced but did not reach the cut off threshold for analysis. It does not appear that sugar metabolism is slowed down in this strain as otherwise we may have expected to see such genes upregulated on Day 4. The observed differences in carbohydrate metabolism may reflect the previously noted differences in the types and copy numbers of genes encoding for maltose and maltotriose transporters between the two strains (40–42).

The upregulation of genes associated with amino acid metabolism is significant as the secondary metabolites associated with flavour such as higher alcohols and esters are produced from the catabolism of amino acids. The gene ontology also revealed differences in metabolism between the two strains. For example, genes associated with methane, butanoate and sulphur metabolism, which can also contribute to flavour profiles, are upregulated in Group II but not in Group I.

Gene ontology analysis confirm the over representation of Se alleles in the upregulated genes in the Group II strain and additionally identified specific pathways where either Sc or Se alleles are exclusively enriched. Considering the reduced Sc content in the Group I strain, it is surprising that Sc alleles appear to be over-represented in genes enriched in several pathways. Specifically, we see Sc alleles contributing to Arginine and Lysine biosynthesis and Tyrosine metabolism at a higher level than expected from the Sc:Se ratio for the whole genome. We also observed that Sc alleles are more likely to be downregulated than Se alleles in both Group I and Group II strains however specific pathways here also contain exclusively Se alleles.

Previous studies in the haploid *S. cerevisiae* showed that the steady state levels of mRNA transcripts are tightly controlled to maintain homeostasis. Disruption of mRNA turnover, for example by deletion of the major 5’ to 3’ exonuclease *XRN1*, is compensated by changes to RNA polymerase II transcription to restore homeostasis (47, 48). Conversely, increased transcription rates resulting from increased gene copy number lead to compensatory changes in mRNA turnover to preserve the expected steady state levels of mRNAs (49). However, a recent analysis of laboratory and wild *S. cerevisiae* strains revealed no evidence of dosage compensation and instead described a direct correlation between gene copy number and gene expression levels (50).

Here we show that the aneuploid nature of the *S. pastorianus* genomes directly contributes to the gene expression patterns during fermentation. We observed a positive correlation between orthologue copy number and the steady state levels of orthologue transcripts in the Group II strain: as the copy number of Sc or Se genes increases so too does the associated levels of transcripts. Interestingly, when the orthologue gene copy number is 1:1, we observed that the Se alleles showed higher levels of transcripts than the Sc alleles in all conditions tested. Surprisingly, the pattern of usage of orthologues is different in the Group I strain. While there is less variation in the ratios of Sc:Se genes with most orthologues present in a 1:2 ratio, here we observed that Sc orthologues produce higher levels of transcripts than the Se counterpart although there is a greater variation in the distribution of gene expression patterns amongst orthologues at this ratio in the Group I strain. At the same ratio in the Group II strain, Se alleles predominate. The over-representation of *S. cerevisiae* alleles in some biochemical processes such as amino acid metabolism may explain the selective retention of *S. cerevisiae* chromosomes in the Group I strain as there may have been a selective evolutionary pressure to retain specific genes required during fermentation.

The *S. eubayanus* parent of the lager yeasts is a cryotolerant strain and thus it might be hypothesised that Se alleles will be favoured during the fermentations which are carried out at 13°C. We did observe a greater contribution of Se alleles to the transcriptome in the Group II strain during fermentation conditions, however, it is not universal, and this contribution is tempered by gene copy number effects. A transcriptome analysis for the Group I strain, CBS1513, had also observed an overrepresentation of *S. eubayanus* when cultures were grown at low temperatures (38).

To compare the overall steady state levels of protein encoding transcripts in the Group I and II strains, we created combined transcriptomes for each strain by summing the Sc and Se transcripts that encode for the same protein. Using this approach, we observed that that overall transcription patterns were similar between the two strains indicating that despite the significant differences at a genome level and pathways of evolution, the two lager yeast groups display similar transcription patterns in fermentation conditions. Nevertheless, gene ontology analysis identified some unique transcription patterns. Consistent with what was observed for the differential expression of Sc and Se alleles within a strain, the gene expression patterns in both strains were significantly influenced by gene copy number.

Thus, our analysis of the expression of gene orthologues within a strain and between strains indicates that the aneuploid genomes of the lager yeasts create complex patterns of gene expression during fermentation and that gene copy number plays a crucial role in the gene expression patterns both within a strain and between strains.

## Materials and Methods

### Yeast strains and growth conditions

The Group I strain CBS 1538 was obtained from the Collection de Levures d’Interet Biotechnologique, Paris, France and Group II strain Weinstephan 34/70 was kindly supplied by Dr. Jurgen Wendland, Geisenheim Hoch Universitat, Germany. For propagation, strains were grown in YPDM medium (1% (w/v) yeast extract, 2% (w/v) peptone and 1 - 2 % (w/v) of both dextrose and maltose at 20-25°C; 2% YPDM contains 1% dextrose and 1% maltose.

For RNA extractions, cells were grown in 2% YPDM overnight and were then washed with sterile water and inoculated at a cell density of 1×10^6^ cells/mL into 60 mL of minimal medium (0.17% Yeast Nitrogen Base w/o amino acids and ammonium salts supplemented with 1% of dextrose and 1% of maltose, and 0.5% (NH4)2SO4 as a nitrogen source). The cultures were grown at 20°C, in triplicate, and cells were harvested in the exponential phase.

Small-scale fermentations (10mL) and large-scale fermentations (2L) were carried out in 10% wort containing 1mM ZnSO_4_ (Spraymalt, Brewferm, Amber 18EBC, Brouland, Belgium) in 15 mL glass test tubes or 3L tall tubes in triplicate or duplicate respectively.

Cells were first propagated in 200 mL of 4% YPDM at 25°C for two days and were pitched at a cell density of 1.5×10^7^/mL. The tubes were fitted with a water trap airlock attached to a bung and incubated at 13°C at a 45° angle without shaking. The specific gravity of the wort was measured at the start of fermentation and at intervals throughout the fermentation using a refractometer (*HANNA*, Romania).

### Flavour Analysis

Analysis of volatile compounds in wort at the end of fermentation were conducted at The Research and Innovation Centre, Edmund Mach Foundation S.Michele all’Adige, Italy as previously described (51). Briefly, 2.5 mL of the samples in 20 mL vials were supplemented with sodium chloride to a final concentration of 40 mg/mL and 25 μL of the 2-octanol as the internal standard (final concentration 200 μg/L). All samples were incubated for 10 min at 40°C, then the volatile compounds were collected on a divinylbenzene / carboxen / polydimethylsiloxane fibre (DVB-CAR-PDMS) coating 50/30 μm, and 2-cm length SPME fibre purchased from Supelco (Sigma Aldrich, Milan, Italy) for 40 min. GC analysis was performed on a Trace GC Ultra gas chromatograph coupled with a TSQ Quantum Tandem mass spectrometer (Thermo Electron Corporation, USA) (51). Identification of compounds was based on comparison with a mass spectral database (NIST version 2.0) and with reference standards when available. The relative amount of each volatile was expressed as μg/L of 2-octanol (52).

### DNA extraction, *de novo* genome sequencing and copy number variation analysis

DNA extraction was carried out following the protocol described by (53). Briefly, yeast cells were grown in an overnight culture of 5 ml of 2% YPDM. Cells were pelleted and washed with distilled water and centrifuged. The pellet was resuspended in 500 μL of 0.9 M sorbitol and 0.1 M EDTA, pH 7.5 and 30 μL of a solution of lyticase (1.5 mg/ mL) was added and incubated for 20-40 min. After this, spheroplasts were recovered centrifuged and resuspended in 500 μL of 50 mM Tris HCl and 20 mM EDTA, pH 7.4 and 13 μL of 10% SDS. Samples were incubated for 65°C for 5 −10 min. 200 μL of 5M potassium acetate were added and tubes were incubated on ice for 10 min. After centrifugation, 700 μL of 100% isopropanol was added to the supernatant, the samples incubated at room temperature for 10 min and then recentrifuged. The pelleted DNA was washed twice with 70% ethanol, air-dried at room temperature, and resuspended in 50 μL of water.

Total DNA was treated with RNAse A (10 μg/mL) for 30 min at 37°C. DNA was precipitated and resuspended in 50 μL of water. The quality of the DNA was determined by agarose gel electrophoresis and by determining the OD_260_:OD_280_ ratio using a Nanodrop spectrophotometer (Epoch 2, Agilent BioTek). *De novo* genome sequencing was carried out by Novogene (www.novogene.com) with Illumina technology on paired-end reads (150bp).

After trimming paired-end reads with trim_galore (version 0.6.6, calling cutadapt version 2.10), the reads were mapped with bowtie2 (version 2.4.2) (54, 55) to the parental genomes of *S. pastorianus* CBS1483, (assembly ASM1102231v1) https://www.ncbi.nlm.nih.gov/genome/?term=Saccharomyces+pastorianus, or to a combined genome of *S. cerevisiae* and *S. eubayanus* derived from *S. cerevisiae* S288C, assembly R64 https://www.ncbi.nlm.nih.gov/genome/?term=Saccharomyces+cerevisiae and *S. eubayanus* (SEU3.0) https://www.ncbi.nlm.nih.gov/genome/?term=Saccharomyces+eubayanus.

In addition to the ‘--fast-local’ presets, parameters were set to limit the insert size to lengths between 200 and 1000 basepairs. Mapped reads were filtered through samtools (version 1.11) to exclude reads that were not mapped in pairs and were not the primary alignment (-F 268) and to include only reads with high mapping quality (-q 44).

For hybrid chromosomes, genes were assigned as *S. cerevisiae* or *S. eubayanus* based on their location relevant to known recombination sites and their % identity to the parental genomes. Reads from *de novo* DNA sequencing, mapped against the combined genomes of the reference strains of *S. cerevisiae* and *S. eubayanus* were transformed into sorted BAM files using samtools and the data was extracted as reads/500 bp and then normalized by the size of the library (total number of reads) to give an estimated chromosome copy number (56).

### RNA extraction and Sequencing

RNA extraction was carried out following the protocol previously described (57) but with some modifications. Cells were resuspended in 400 μL of buffer containing 50 mM of NaOAc, pH 5.2, 10 mM EDTA pH 8.0 and 40 μL of 10% SDS, 400 μL of Phenol-Chloroform-Isoamyl alcohol (PCA; 25:24:10), pH 4.2, and 100 μL of acid washed glass beads were added. The tubes were placed in a heat block at 65°C for 10 min and vortexed intermittently. The samples were placed on ice for 5 min. After centrifugation, the supernatant was re-extracted with PCA, pH 4.2 and then once with chloroform. Nucleic acids in the supernatant were precipitated, washed twice with 70% ethanol and then resuspended in 100 μL RNase-free H_2_O. The RNA concentration was measured using a Nanodrop spectrophotometer and RNA integrity was checked by Agilent 2100 Bioanalyzer as per manufacturer’s instructions.

RNA sequencing was conducted on cDNA libraries using Illumina technology at the Genomic Technologies Core Facility at the University of Manchester. As a reference genome for the mapping, the combined *S. cerevis*iae and *S. eubayanus* genomes were used but with variations as present in the re-sequenced CBS1538 and WS 34/70 strains. For that, consensus Fasta sequences were generated from the WGS data mapped to *S. cerevis*iae and *S. eubayanus* through an in-house Perl script. To transfer gene annotation from those genomes, insertions and deletions were ignored and only nucleotide replacements were taken into account. Approximately 88k and 126k reads were unmapped for CBS1538 for WS 34/70 respectively. The paired RNA-Seq data were mapped against these strain-specific reference genomes using STAR (58). Multimapping reads are dealt with through the outSAMmultNmax parameter set to 1 and outMultimapperOrder set to ‘random’. Additionally, only the highest quality alignments were kept by filtering with samtools (-q 255). To aggregate reads counts, the tool “featureCounts” was used on the mapped data together with the gene annotation from the *S. cerevis*iae and *S. eubayanus* genome (59). With the −B and the −P options set, only read counts that have both ends aligned within a distance of 50-600 basepairs were considered.

Read counts from the RNA mapping were uploaded into iDEP9.1 (60). Data were transformed for clustering and PCA by EdgeR using a minimum CPM of 0.5 in 1 library and adding a pseudocount of 4. PCA analysis was used to check the reproducibility of replicates. Outlier replicates were removed and data reprocessed. Differentially Expressed Genes (DEG) were calculated using DESeq2 with a False Discovery Rate (FDR) cutoff of 0.05 and a minimum log_2_ fold change of ≥1 or ≤-1. Venn diagrams were made using the online tool (http://bioinformatics.psb.ugent.be/). The copy number for each gene was extracted from the reads coverage of the *de novo* genome sequencing of the *S. pastorianus* strains.

Enrichment of DEG was carried out using ClueGO (61). Parameters used on ClueGo were set as follows: *Saccharomyces cerevisiae* S228C was used as a Load Maker List. KEGG Ontology was used for the analysis with the following filters, p-value less than 0.05, minimum number of four genes per pathway, pathways containing more than 25% of the genes regulated, Kappa score was set at 0.4. A two-sided hypergeometric test and a Benjamini-Hochberg pV correction was applied. *S. eubayanus-like* allele names were translated into *S. cerevisiae* gene names for ClueGo analysis.

A combined transcriptome of *S. pastorianus* genes was generated by summing the reads for *S. cerevisiae* and *S. eubayanus* orthologues using the consolidate function in Excel. Unmatched genes were retained in the combined transcriptome with reads mapped respectively to the *S. cerevisiae* or *S. eubayanus* reference genomes. The combined genome contained 7568 unique gene ids. The copy number for each gene was extracted from the reads coverage of the *de novo* genome sequencing of the *S. pastorianus* strains and were summed for the matched orthologues to give a total gene copy number for each gene id.

### Statistics

False discoveries rates for DEG and gene ontologies were determined using the Benjamini-Hochberg correction. Chi-squared tests were used to determine the enrichment of genes from sub-genomes under the different physiological conditions used.

## Acknowledgements

We thank Penghan Zhang, Silvia Carlin and Urska Vrhovsek for assistance with GC/MS analysis.

## Notes

### Competing Interest Statement

The authors have declared no competing interest.

